# Fractional Laser Treatment Boosts Systemic Anti-Tumor Immunity in Combination with OX40 Agonist and PD-1 Blockade

**DOI:** 10.1101/2021.11.29.470433

**Authors:** Masayoshi Kawakubo, Joshua Glahn, Zhipeng Tao, Shadmehr Demehri, Dieter Manstein

## Abstract

While ablative fractional photothermolysis (aFP) with a 10,600 nm CO_2_ laser is employed for a wide variety of dermatologic conditions, its applications in oncology are relatively unexplored. Building off our previous work, we investigated the effect of unilateral aFP treatment in combination with anti-PD-1 blocking antibody and OX40 agonist on bilateral tumor growth and remission. A CT26 wild type (CT26WT) colon carcinoma cell line was established bilaterally on the hind flanks of a standardized mouse model and tumor characteristics were investigated on aFP treated and untreated sides. Remarkably, triple therapy with fractional CO_2_ laser in combination with anti-PD-1 antibodies and OX40 agonists resulted in significantly slower tumor growth and complete remissions on bilateral tumors. Flow cytometric analysis showed the triple treatments elicited an increase of granzyme B+ CD8+T cells due to synergistic effect of aFP treatment and the checkpoint molecules, including the induction of CD103+ CCR7+ dendritic cells (DCs) in aFP-treated tumor by aFP treatment, XCR1+ DCs in drainage lymph node by anti-PD-1 inhibitor and OX40+ Ki67+ CD8+ T cells in the lymph node by OX40 agonist. Triple therapy-mediated tumor regression and survival was abrogated upon CD8+ T cell depletion. Importantly, when two mismatched cancer cells were implanted into mice, the effect of the triple therapy on distant tumor was abrogated, showing antigen specificity of the T cell immunity induced by triple therapy. This study highlights the efficacy of aFP a novel adjuvant for current cancer immunotherapeutics.

## Introduction

Ablative fractional photothermolysis (aFP) is a laser treatment modality used to generate a regular pattern of microscopic zones of thermal damage in tissue (1). aFP is used for a range of dermatologic conditions including treatment of photodamaged skin, dyschromia, rhytids, and revision of acne, surgical, and burn scars (1–5). The therapy consists of a fractionated pattern of microscopic treatment zones (MTZ) created by focused laser beams at wavelengths highly absorbed by molecules in the tissue. These MTZs consisted of an empty column of ablated tissue surrounded by a ring of denatured collagen and thermally damaged tissue (6, 7). MTZ usually measure less than 0.5 mm across, however, characteristic such as diameter and depth depend on adjustable laser parameters. aFP typically damages only a small fraction of the tissue (often between 5-20%), leaving the majority of tissue unharmed (8). By creating a path through the stratum corneum, aFP has been used to facilitate transcutaneous drug delivery of chemotherapeutics and photosensitizers in the treatment of cutaneous cancers (9). However, the use of aFP as a primary treatment modality remains poorly understood.

In a previous study we investigated the role of aFP in inducing anti-tumor immunity to promote tumor regression both locally and systemically (10, 11). We demonstrated that, in combination with an anti-PD-1 inhibitor, aFP promotes the proliferation of tumor antigen specific CD8+ T-cells in both treated and untreated tumors in a bilateral mouse tumor model. While treatment led to near complete remission in irradiated tumors, aFP+ anti-PD-1 inhibitor therapy only resulted in contralateral tumor size and growth rate reduction in 33% of mice (11). We theorize that the limited success of systemic anti-tumor immunity despite bilateral infiltration of the CD8+ T-cells is the ratio of regulatory T-cells (T_reg_) to CD8+ T-cells in the contralateral tumor, blocking the antigen-specific immunity promoted by aFP.

To overcome this proposed immune blockade, an OX40 agonist was added to aFP + anti-PD-1 inhibitor therapy. OX40 agonists are shown to promote the activation, proliferation and effector function of killer CD8+ T-cells (12–14) in addition to promoting the proliferation and effector function of helper CD4+ T-cells (15–17) and depleting the number of T_reg_ cells (18–21). By repressing the regulatory checkpoints inhibiting CD8+ T-cell effects systemically, triple therapy with aFP+ anti-PD-1 inhibitor + OX40 agonist has the potential to become a novel immunomodulatory therapy for promoting tumor regression both at the treatment site and in remote tumors not treated with aFP.

## Results

### aFP treatment boosts effect of systemic anti-tumor immunity induced by anti-PD-1 inhibitor and OX40 agonist combination therap

We established a two-tumor mouse model, with identical tumors seeded on each hind leg, to evaluate the effect of aFP treatment in conjunction with immune checkpoint molecules on systemic induction of anti-tumor immunity. In the initial experiment, tumors were left for 6 days post inoculation, reaching an average diameter of 4mm, before aFP treatment. Intraperitoneal treatment with anti-PD-1 inhibitor and/or OX40 agonist occurred on days 6, 8, 10, 12, and 14 after inoculation. Triple therapy with aFP + anti-PD-1 + OX40 agonist led to a significant reduction in tumor volume and growth rate following treatment (Figure 1a and b). Triple therapy resulted in complete tumor remission in all mice with uniform survival past the 90-day predetermined endpoint (Figure 1c). Significance value between survival curves are shown in Table 1. Of note, there was no significant difference between triple therapy and therapy with anti-PD-1 + OX40 agonist alone. To test the improved efficacy of triple therapy over double therapy, a second round of experiments was conducted with treatment beginning 12 days after inoculation and an average tumor diameter of 7mm. aFP treatment was performed on day 12 and anti-PD-1 inhibitor and/or OX40 agonist were administered on days 12, 14, 16, 18, and 20 after inoculation. Triple therapy in the delayed treatment group led to a significant reduction in tumor volume and growth rate over therapy with anti-PD-1 inhibitor and OX40 agonist (Figure 2a, b and c). Both aFP treated and contralateral tumors achieved complete remission in the triple therapy group and 8 of 10 mice survive to 90 days compared to tumor shrinkage in 3 of 10 mice in the double therapy group (Figure 2 d). Significance values for differences between survival curves are shown in Table 2.

**Figure 1.**
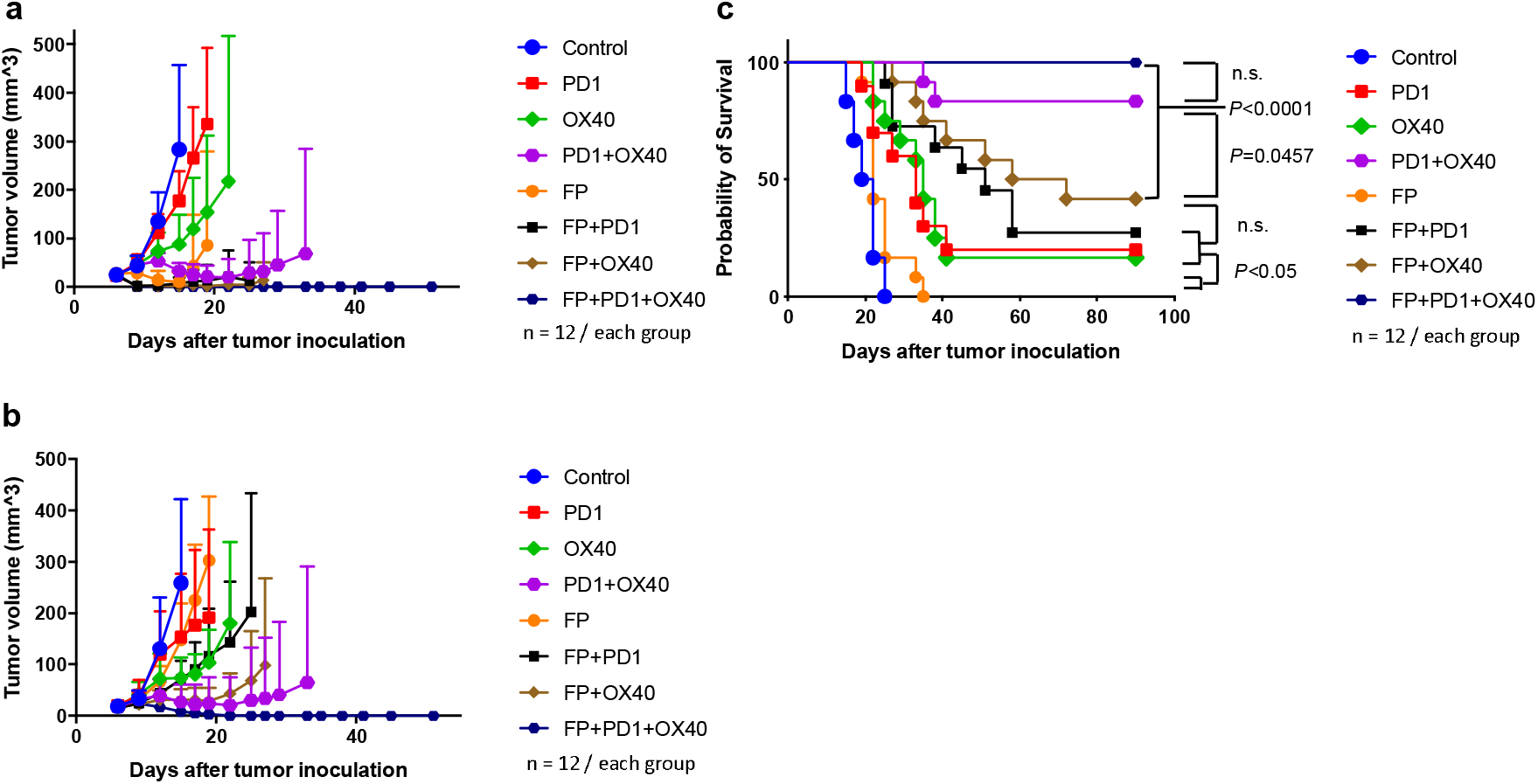
6-day inoculation tumor volume and survival curves post treatment. To investigate whether aFP treatment can induce systemic anti-tumor immunity, we established a mouse model with tumors on both hind legs. Tumors on the left were treated with aFP and growth of both tumors was observed. aFP laser irradiation was performed 6 days after tumor inoculation. Anti-PD-1 inhibitor and OX40 agonist were administered intraperitoneally at a dose of 200 μg per mouse on days 6, 8, 10, 12 and 14 after tumor cell inoculation. (a) Average tumor volume curves on the aFP-treated legs of mice treated on day 6. (b) Average tumor volume curves on the untreated contralateral legs of mice treated on day 6. (c) Kaplan-Meier survival curves of mice receiving tumor inoculation, aFP-treated on day 6. The significance values for the difference between the survival curves are shown in Table 1. : Control vs. atni-PD-1: *P* = 0.0007, Control vs. OX40: *P* < 0.0001, Control vs. Anti-PD-1 + OX40: *P* < 0.0001, Control vs. aFP: *P* = 0.0217, Control vs. aFP + anti-PD-1: *P* < 0.0001, Control vs. aFP + OX40: *P* < 0.0001, Control vs. aFP + anti-PD-1 + OX40: *P* < 0.0001, anti-PD-1 vs. OX40: n.s., anti-PD-1 vs. Anti-PD-1 + OX40: *P* = 0.001, anti-PD-1 vs. aFP: *P* = 0.0281, anti-PD-1 vs. aFP + anti-PD-1: n.s., anti-PD-1 vs. aFP + OX40: n.s., anti-PD-1 vs. aFP + anti-PD-1 + OX40: *P* < 0.0001, OX40 vs. Anti-PD-1 + OX40: *P* = 0.0007, OX40 vs. aFP: *P* = 0.0018, OX40 vs. aFP + anti-PD-1: n.s., OX40 vs. aFP + OX40: n.s., OX40 vs. aFP + anti-PD-1 + OX40: *P* < 0.0001, Anti-PD-1 + OX40 vs. aFP: *P* < 0.0001, Anti-PD-1 + OX40 vs. aFP + anti-PD-1: *P* = 0.01, Anti-PD-1 + OX40 vs. aFP + OX40: *P* = 0.0457, Anti-PD-1 + OX40 vs. aFP + anti-PD-1 + OX40: n.s., aFP vs. aFP + anti-PD-1: *P* < 0.0001, aFP vs. aFP + OX40: *P* < 0.0001, aFP vs. aFP + anti-PD-1 + OX40: *P* < 0.0001, aFP + anti-PD-1 vs. aFP + OX40: n.s., aFP + anti-PD-1 vs. aFP + anti-PD-1 + OX40: *P* = 0.0002, aFP + OX40 vs. aFP + anti-PD-1 + OX40: *P* < 0.0001.

**Figure 2.**
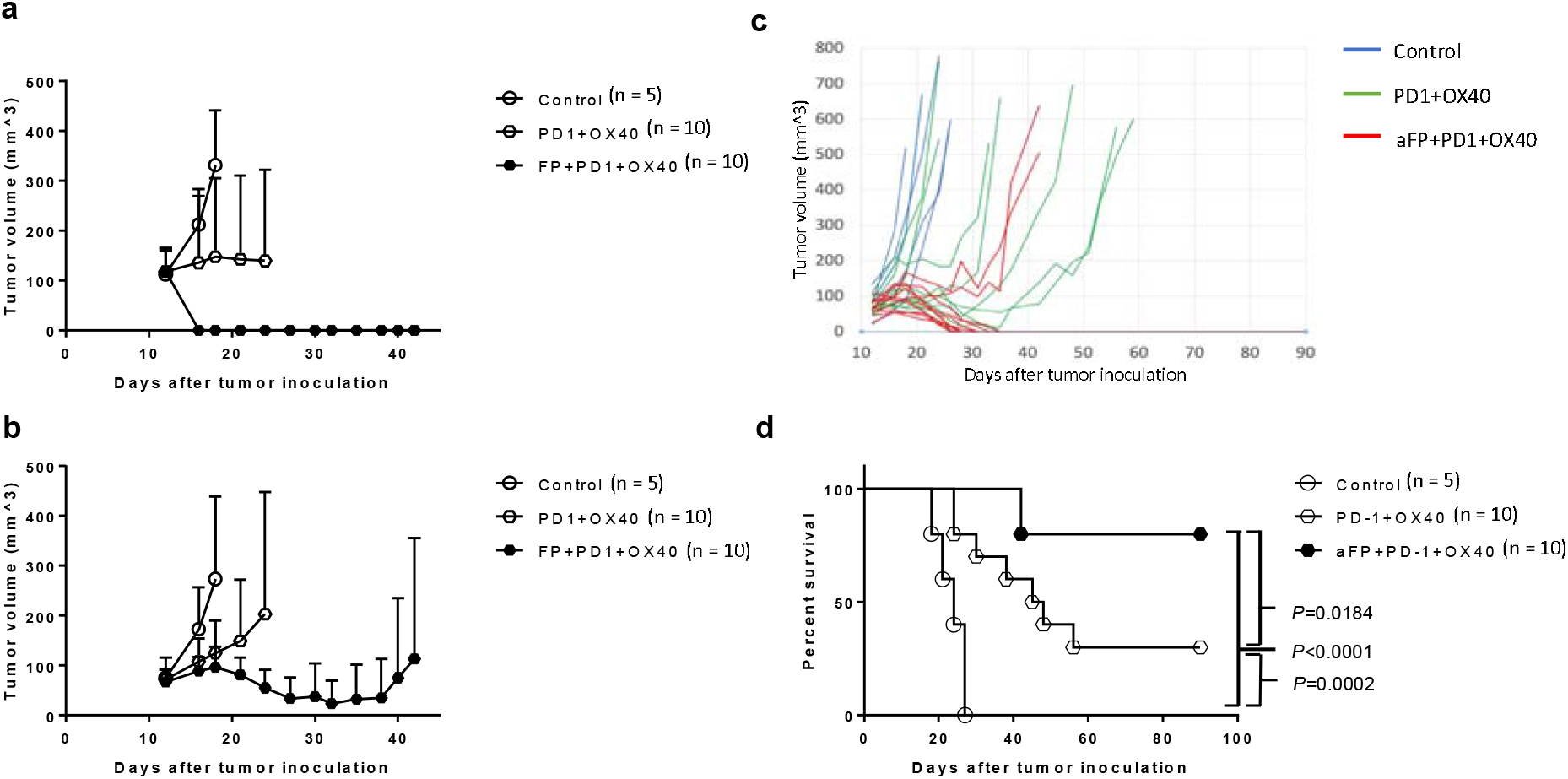
12-day inoculation tumor volume and survival curves post treatment. To investigate the relative efficacy of anti-PD-1 + OX40 agonist therapy vs triple therapy (aFP + anti-PD-1 + OX40 agonist) in large tumors, aFP and immunomodulatory treatment was started 12 days after tumor inoculation. Anti-PD-1 inhibitor and OX40 agonist were administered intraperitoneally at a dose of 200 μg per mouse on days 12, 14, 16, 18 and 20 after tumor cell inoculation. (a) Average tumor volume curves in tumors treated with aFP on day 12. (b) Average tumor volume curves on the untreated contralateral tumors in mice treated with aFP on day 12. (c) Individual tumor volume curves on the untreated contralateral tumor in mice treated with aFP on day 12. (d) Kaplan-Meier survival curves in mice receiving aFP treatment 12 days post tumor inoculation. Significance values between survival curves are: Control vs. Anti-PD-1 + OX40: *P* = 0.0022, Control vs. aFP + anti-PD-1 + OX40: *P* < 0.0001, Anti-PD-1 + OX40 vs. aFP + anti-PD-1 + OX40: *P* = 0.0252.

### Surviving mice develop long-term anti-tumor immunity

To evaluate for long-term induction of anti-tumor immunity, mice from the double and triple therapy groups that survived 90 days post original inoculation with complete tumor remission and a group of age-matched tumor naive mice received a subcutaneous inoculation of 3.5 × 10^5^ CT26WT cells in right thigh (aFP-untreated thigh in case of aFP-treated mice). While tumors on the tumor-naive control mice progressed over time leading to no survivors at 30 days, tumors in the rechallenged double and triple therapy groups showed no signs of tumor progression and remained tumor–free for at least 60 days post inoculation (Figure 3). Significance values between the survival curves are: control vs. anti-PD-1 + anti-OX40 (*p*<0.0001), control vs. aFP + anti-PD-1 + anti-OX40 (*p*<0.0001).

**Figure 3.**
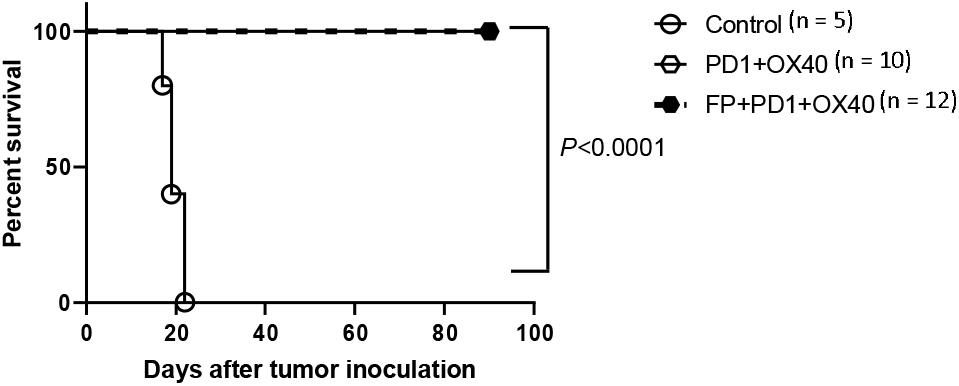
Survival curves on rechallenge. Kaplan-Meier survival curves of mice receiving the rechallenge test with CT26WT cells. Each mouse was inoculated subcutaneously with 3.5 × 10^5^ CT26WT cells on the aFP-untreated contralateral thigh. Significance values between survival curves are: control mice vs. survival mice in anti-PD-1 + OX40 and aFP + anti-PD-1 + OX40: *P* < 0.0001.

### The triple therapy induces tumor-infiltrating CD8+ and granzyme B+ CD8+ T lymphocytes in aFP-untreated contralateral tumors

To investigate the mechanism by which triple therapy led to regression in untreated contralateral tumors, flow cytometry of untreated tumor cells 5 days post-aFP were analyzed for the concentration of tumor-infiltrating CD3+, CD4+, CD8+, and granzyme-B+ CD8+ T-cells, along with Foxp3+ regulatory T-cells (T_regs_). CD8+ and granzyme B+ CD8+ T-cells significantly increased by tumor weight in the triple therapy group compared to all other treatment groups (CD8/weight: control, anti-PD-1, anti-OX40, aFP, aFP + anti-PD-1 or aFP + anti-OX40 vs. aFP + anti-PD-1 + anti-OX40 (*p* < 0.05; Figure 4d-e). However, there was no significant difference in CD3+ and CD4+ T-cell concentrations between experimental groups (Figure 4a-b). Furthermore, there was no significant difference between number of T_reg_ cells between groups (Figure 4c), resulting in an increased ratio of CD8+ T-cells to T_reg_ in triple therapy (control, anti-PD-1, or aFP + anti-PD-1 vs. aFP + anti-PD-1 + anti-OX40 (*p* < 0.05; Figure 4f).

**Figure 4.**
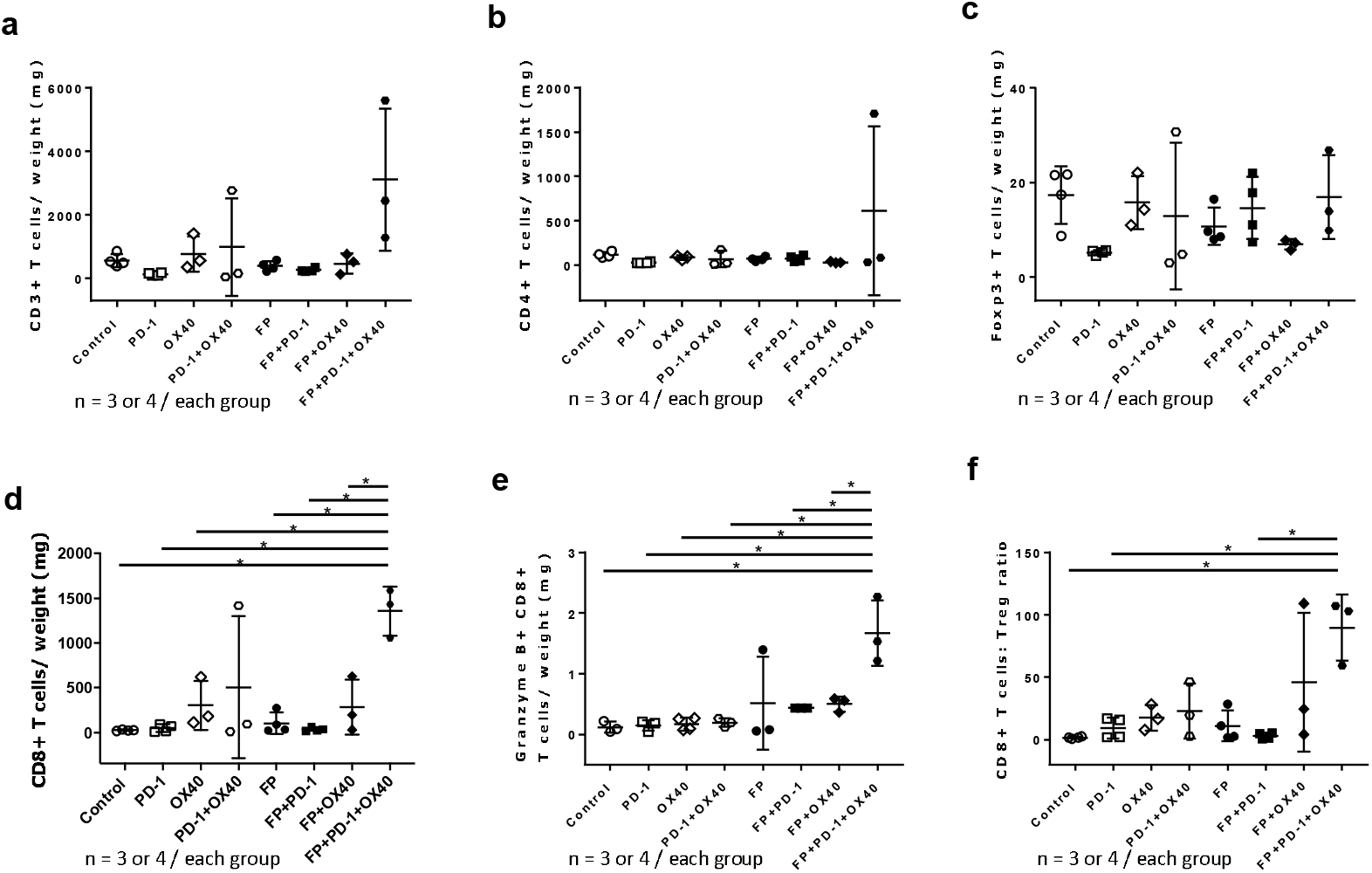
Flow cytometric analysis of tumor infiltrating lymphocytes 5 days post aFP treatment in untreated contralateral tumors. Flow cytometry analysis was performed 5 days after aFP treatment to investigate the number of CD3+, CD4+, and CD8+, and granzyme B+ T-cells in addition to the ratio of T_regs_ expressing CD4 and Foxp3 in the untreated contralateral tumor. (a) Proportion of CD3+ T-cells cells normalized to tumor weight. (b) Proportion of CD4+ T-cells normalized to tumor weight. (c) Proportion of Treg normalized to tumor weight. (d) Proportion of CD8+ T cells normalized to tumor weight. (e) Proportion of granzyme B+ CD8+ T cells normalized to tumor weight. (f) Ratio of CD8+ T cells to Tregs (CD4+Foxp3+).

### aFP + anti-PD-1 therapy increases CD103+ CCR7+ dendritic cells in aFP-treated tumor and XCR1+ DCs in drainage lymph node

To investigate how triple therapy significantly increases tumor-infiltrating CD8+ T lymphocyte migration into contralateral tumors, we measured the number of CD45+ CD11c+ MHC II+ dendritic cells (DCs) present in in aFP-treated tumor and associated drainage lymph nodes 3 days post-treatment using flow cytometry. For this subset of experiments, we did not used OX40 agonists due to the lack of OX40 receptors on target DCs. Initially, we measured the concentration of CD103+ CCR7+ dendritic cells in aFP-treated and control tumors. CD103+ DCs were chosen as the specific subpopulation with the ability to induce proliferation of naïve CD8 T-cells (22) in the presence of antigen, and migration of DC to local lymph nodes is directly mediated by the CCR7 chemokine receptor (responsive to chemoattractant CCL19 and CCL21) (24). We found an increase in the concentration of CD103+ CCR7+ DCs normalized to tumor weight in aFP treatment groups (aFP and aFP + anti-PD1) when compared to control and anti-PD1 only groups (Figure 5a), with the aFP + anti-PD-1 group showing a significant increase in DC concentration over anti-PD-1 alone (*p* < 0.05).

**Figure 5.**
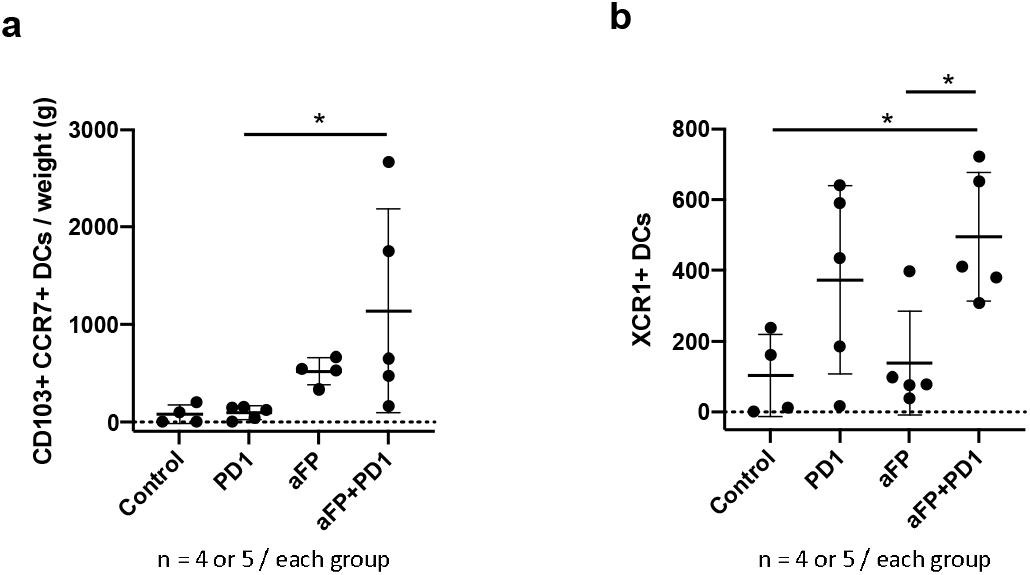
Flow cytometric analysis of dendritic cells (DCs) 3 days post aFP treatment in aFP-treated tumor and local drainage lymph node. Flow cytometry analysis was performed 3 days after aFP treatment to investigate the number of CD103+ CCR7+ DCs in aFP-treated tumor and XCR1+ DCs in drainage lymph node on the treated side. (a) Proportion of CD103+ CCR7+ DCs normalized to tumor weight. (b) Absolute number of XCR1+ DCs in drainage lymph node.

Next, we measured the number of DCs expressing XCR1 in drainage lymph node. The interaction between naive CD8+ T-cells and cross-presenting DCs is mediated by the XCR1-XCL1 axis. XCL1 is a chemoattractant secreted by CD8+ T-cells and is sensed by XCR1 ligands on DCs. This interaction enhances cross-priming, making the XCR1-XCL1 axis an integral component in the development of efficient cytotoxic immunity (25, 26). As shown in Figure 5b, XCR1+ DC concentrations increased significantly in the aFP + anti-PD-1 group as compared to control and aFP only groups (*p* < 0.05). Taken together, these data suggest that aFP treatment recruits DCs with the capability of inducing proliferation of naive CD8+ T-cells, enabling their migration to drainage lymph node on the aFP-treated tumor side, and anti-PD-1 inhibitor increases DCs and is an integral component in the development of efficient cytotoxic immunity.

### Triple therapy induces expression of OX40 and PD-1 on CD8+ T cells in local drainage lymph nodes

To investigate why triple therapy resulted in higher concentrations of tumor-infiltrating CD8+ T lymphocytes in the contralateral, untreated tumor, we quantified expression of PD-1, OX40, and Ki67 on CD8+ T-cells in local lymph nodes 5 days after aFP treatment. OX40 and PD-1 are expressed rapidly after antigen stimulation on CD8+ T-cells (27, 28), meaning the expression of OX40 and PD-1 is a marker of stimulation.

Flow cytometry confirmed the expression of OX40+ PD-1+ CD8+ T-cells in drainage lymph nodes on the aFP-treated side with groups receiving triple therapy displaying significantly higher number of T-cells. The basis of immunomodulation in triple therapy is the ability of OX40 agonists to promote proliferation of OX40-expressing T-cells while the anti-PD-1 antibodies counteract the immune suppression caused by the PD-1/PD-L1 pathway. Significantly increased proliferation of OX40+ CD8+ T-cells in the triple therapy treatment group was confirmed by staining for Ki67, a common marker of proliferation. The experimental groups receiving aFP, OX40 agonists, and PD-1 antagonist (triple therapy) demonstrated significantly higher proportion of Ki67+ staining cells than OX40 agonist alone group (*p* < 0.01) and than control and anti-PD-1 groups (*p* < 0.05; Figure 6a). Moreover, the increase in PD-1+ OX40+ Ki67+ CD8+ T-cells in the triple therapy group increased significantly when compared with any other experimental group except FP+OX40 group, indicating that triple therapy is an effective promoter of antigen-stimulated CD8+ T-cell proliferation, leading to higher systemic concentrations of immune cells and infiltration into contralateral tumors (control, anti-PD-1, anti-OX40, or aFP vs. triple therapy (*p* < 0.005), anti-PD-1 + anti-OX40 or aFP + anti-PD-1 vs. aFP + anti-PD-1 + anti-OX40 (*p* < 0.05); Fig. 6b).

**Figure 6.**
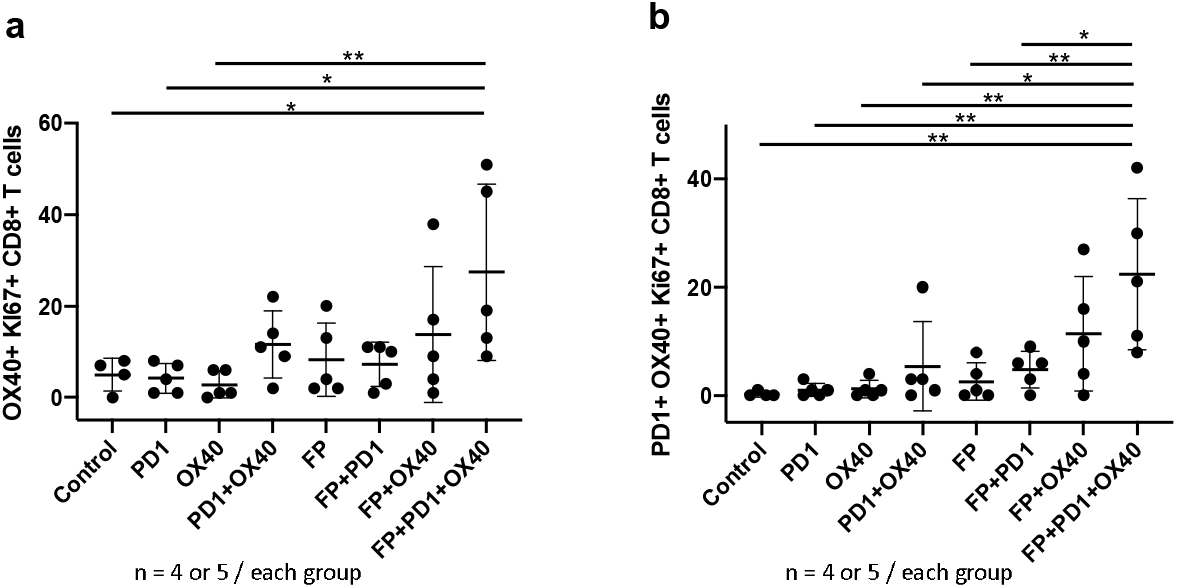
Flow cytometric analysis for OX40+ Ki67+ CD8+ T cells 5 days after aFP treatment in local drainage lymph node. Flow cytometric analysis was performed 5 days post aFP treatment to quantify the number of OX40+ Ki67+ CD8+ T-cells in drainage lymph node on the aFP-treated side. (a) Absolute number of OX40+ Ki67+ CD8+ T-cells in drainage lymph node on aFP-treated side. (b) Absolute number of PD-1+ OX40+ Ki67+ CD8+ T-cells in drainage lymph node on aFP-treated side.

### Adaptive immunity is crucial to eradicating cancer cells after the triple therapy

To confirm the role of adaptive immunity in eradicating cancer cells systemically with triple therapy, both groups received triple therapy and the experimental group received anti-CD8 depletion antibodies to eliminate CD8+ T-cell populations in treated mice. The beneficial effects of triple therapy were counteracted by suppression of CD8+ T-cell function, with the group receiving anti-CD8+ depletion antibodies showing no survival past 30 days and 4 out of 5 mice in the triple therapy control group surviving the full 90 days (*p*=0.0018; Figure 7a-b). These results indicate that CD8+ T-cells and adaptive immunity play a critical role in the therapeutic benefits of triple therapy.

**Figure 7.**
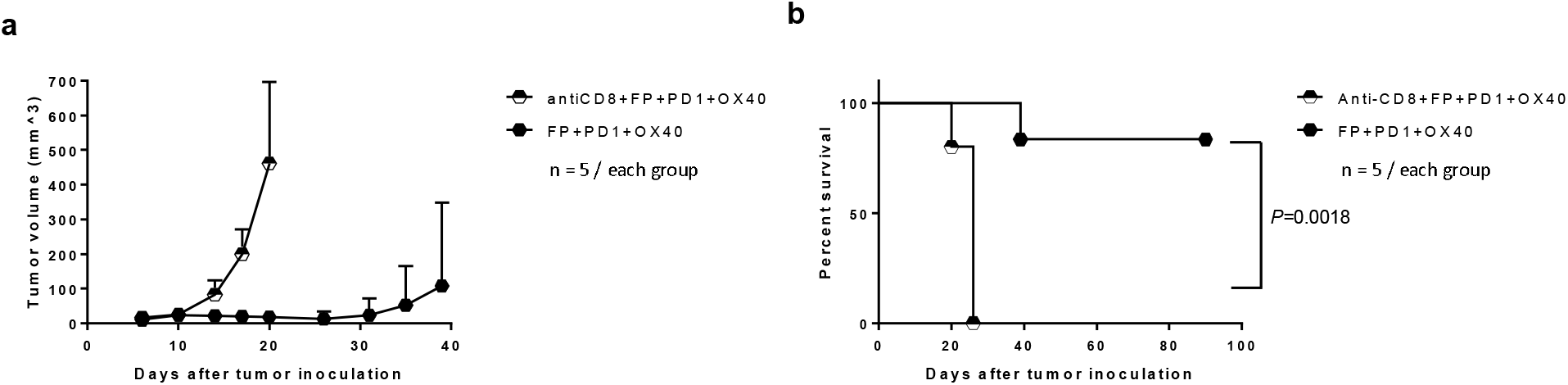
Tumor volume and survival curves after triple therapy with anti-CD8 depletion antibodies. To investigate whether adaptive immunity is necessary to eradicate cancer cells after the triple therapy, tumor inoculation was performed with an anti-CD8 depletion antibody to wipe out CD8+ T-cells in the mouse tumor model. Anti-CD8 depletion antibodies were administered intraperitoneally at a dose of 200 μg every 3 days from the day before tumor inoculation to removal of mice as an endpoint. (a) Difference in tumor volume over time between experimental groups. (b) Kaplan-Meier survival curves showing differences in mortality in mice receiving triple therapy +/- anti-CD8 depletion antibody. The significance values for the difference between the survival curves are: anti-CD8+aFP vs. aFP + anti-PD-1 + OX40: *P* =0.0018.

### Antigen specificity is necessary for systemic anti-tumor immunity induced by aFP

To demonstrate the antigen specificity of triple therapy on distant tumors, we developed a mismatched mouse model by inoculating CT26WT tumor cells in the left leg and 4T1 tumor cells (murine mammary carcinoma) in the right leg. This experimental group was compared with a matched mouse model with inoculation of CT26WT tumor cells in both left and right legs. Triple therapy with aFP was performed on the left haunch only. While the matched mouse group saw complete remission of both tumors and survival of all mice past 90 days, the mismatched mouse group saw complete remission of the aFP treated tumor without a comparable effect on the mismatched 4T1 side, resulting in no survival past 30 days (*p*=0.0035; Figure 8a and b). The benefits did not extend to the contralateral side, suggesting an antigen specific mediated reaction to aFP.

**Figure 8.**
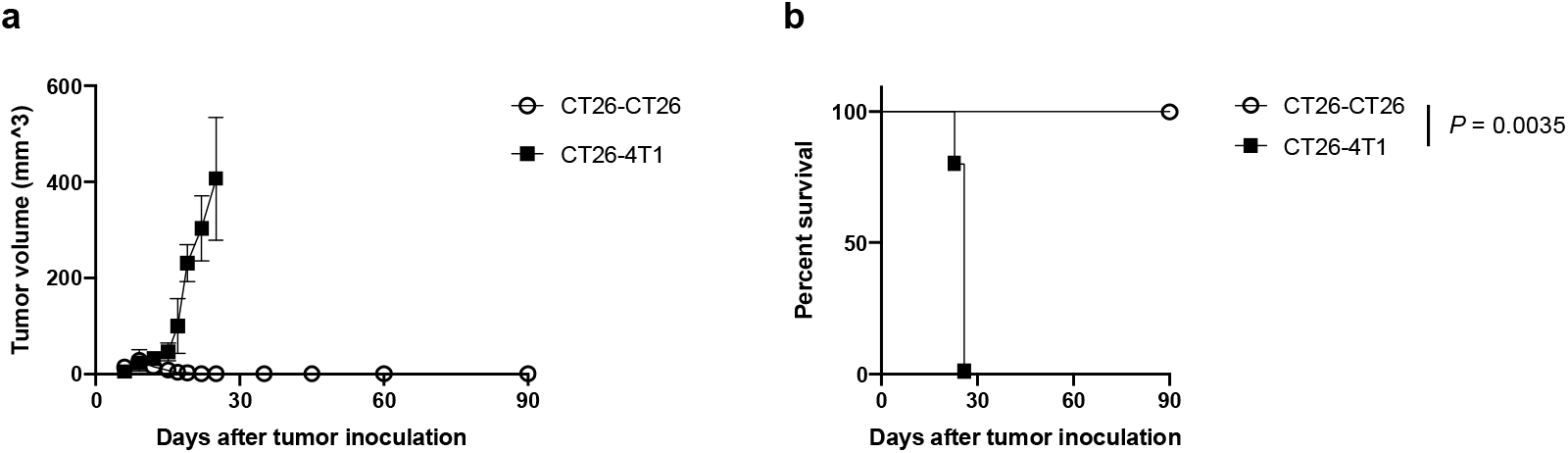
Tumor volume and survival curves after treatment in a mismatched CT26WT and 4T1 tumor model. To determine whether the observed shrinkage of the contralateral, untreated tumor in triple therapy is due to antigen specific immunity induced by aFP, we compared outcomes in matched and mismatched tumor models. In the matched group, mice were inoculated with CT26WT in both legs. In the mismatched group, mice were injected with CT26WT tumor cells in the left leg and 4T1 (murine mammary carcinoma) cells in the right leg. Both groups were treated with triple therapy and aFP was performed on the left CT26WT leg alone. (a) Volume of the right untreated contralateral CT26WT or 4T1 tumor in triple therapy. (b) Kaplan-Meier curves in matched and mismatched tumor model receiving triple therapy. Bars represent SD. The significance values for the difference between the survival curves are: CT26-CT26 vs. CT26-4T1:*P* =0.0035.

### aFP treatment induces expression of HSP70, -90, and calreticulin on tumor cells

There is evidence that extracellular localized and membrane-bound heat shock proteins (HSPs) play a key role in eliciting antitumor immune responses by acting as carriers for tumor-derived immunogenic peptides, enhancing presentation of antigens as targets for the innate immune system (29). Additionally, the expression of calreticulin (CALR), the so called “eat me sign”, has been associated with enhanced immune function and shares a common receptor (LRP1) with HSP70 and -90 on CD103+ dendritic cells (30). To investigate the effect of aFP on transcription of HSP70 and -90 precursors, RNA sequencing was performed 24-hours after tumor irradiation. Both HSPa1a and HSPa1b, mRNA precursors of HSP70, were found to be upregulated in aFP treated tumors as compared to control (Supplemental Table 1). To measure the expression of HSP70, -90, and calreticulin in aFP treated tumor cells, CT26WT colon carcinoma cells were transfected with green fluorescence protein (GFP) for detection. On day 3 after irradiation, flow cytometry confirmed that expression of HSP70, -90, and CALR on GFP transfected cells was significantly higher than in the control group (Figure 9a-c). Furthermore, the proportion of CD103+ DCs expressing LRP1 by tumor weight was significantly higher in the aFP-treated tumor (Figure 9d). The increase in HSP70, -90, and CALR, along with the augmented concentration of their receptor, suggests aFP itself induces expression of molecules which potentially promote anti-tumor immunity.

**Figure 9.**
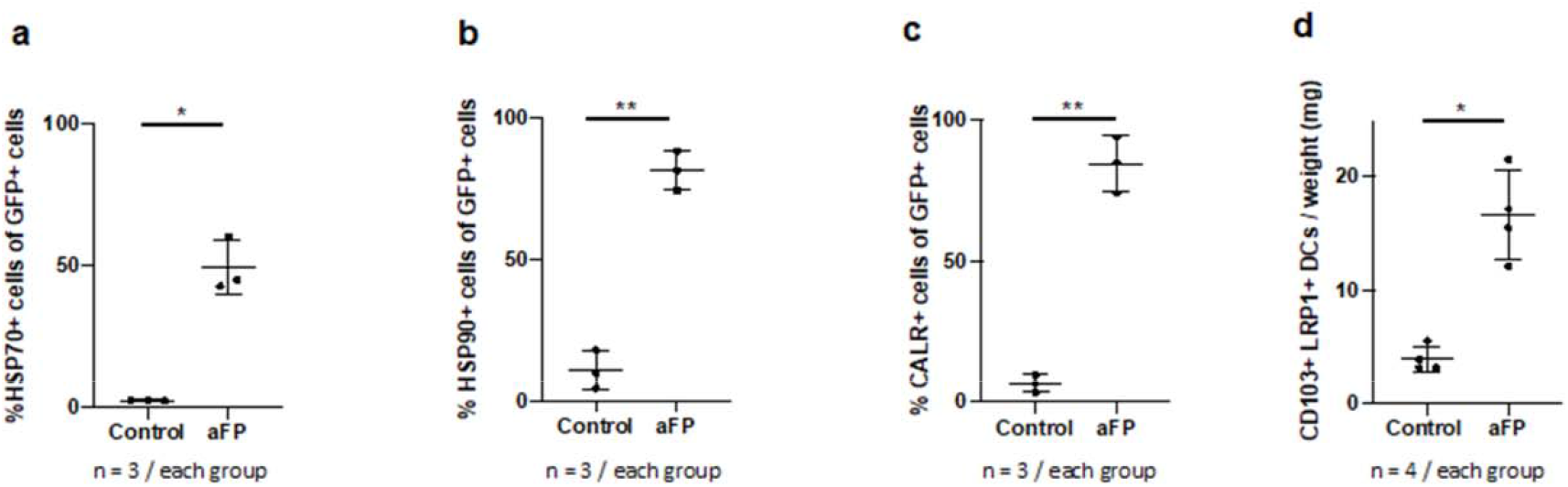
Flow cytometric analysis of DAMPs-expressing tumor cells and DCs expressing LRP1 3 days after aFP treatment in aFP-treated tumors. Flow cytometry analysis was performed 3 days after aFP treatment to investigate percentage of HSP70, HSP90 and CALR-expressing GFP+ tumor cells and the number of LRP1+ DCs in aFP-treated tumors. (a) Percentage of GFP+ cells expressing HSP70. (b) Percentage of GFP+ cells expressing HSP90. (c) Percentage of GFP+ cells expressing CALR. (d) Concentration of LRP1+ DCs normalized to tumor weight.

## Discussion

This study illustrates the benefits of triple therapy with ablative fractional photothermolysis (aFP), anti-PD-1 antibodies, and OX40 agonists in initiating a systemic, tumor specific immune response leading to a significant reduction of tumor burden. While aFP is currently used for a variety of clinical indication, its therapeutic value is generally thought of in terms of localized tissue effects within a limited treatment area for the stimulation of wound healing or tissue regeneration. This study builds on our prior work, expanding the applications of aFP into the world of immunotherapeutics (10, 11). Fractional laser therapy’s ability to induce tumor regression relates to tumor associated antigen (TAA). Although Mroz *et al* demonstrated the utility of TAA in cancer therapy to produce anti-tumor immune activity, antigen expression on tumor cells at baseline is not sufficient to stimulate an immune response to prevent tumor growth (31, 32). The basis of aFP treatment is the release of TAA through thermal denaturation of tumor cells, potentially enhancing anti-tumor immunity by allowing cytotoxic T-cells to recognize and bind the immunodominant peptide epitope (10, 11). The choice of a fractionated laser pattern of approximately 5% microscopic treatment zone (MTZ) to tumor ratio was targeted at enhancing inflammation while avoiding direct killing of the local tumor (Supplement figure1) (10, 11). In our prior study, we found that aFP treatment of CT26WT tumors both induced anti-tumor immunity and improved long term survival (11). Huang *et al* demonstrated that the CT26WT colon carcinoma cells line have a dominant TAA in a single peptide known as AH-1, a non-mutated nonamer derived from the envelope protein (gp70) of an endogenous ecotropic murine leukemia provirus (33). Pentamer staining in our previous study revealed an increase in AH1 specific CD8+ T-cells normalized to tumor weight in aFP-treated groups, supporting the theory that aFP promotes the release of TAA and induces anti-tumor immunity (11). Our current study further substantiates this claim by observing an increase in CD103+ CCR7+ DCs in aFP-treated tumors consistent with the increase in local TAA and the generation of damage-associated molecular patterns (DAMPs) (34). The choice of ideal laser dosimetry, including variation in pulse energy, ablative wavelength, and pattern density to maximize thermal damage and the release of DAMPs deserves further study. The second component of triple therapy, anti-PD-1 inhibitory antibodies, also contributes to anti-tumor immunity. PD-1 is a cell surface protein used to dampen the immune response and is observed in multiple subset of activated lymphocytes, including T- (35), B- (36), and NK-cells (37). PD-1 expression on CD8+ T cells is primarily due to antigen-driven T-cell receptor (TCR) signaling, with expression of the protein within 2-4 hours after antigen exposure and before cell division (27). PD-L1 and PD-L2 are co-inhibitory factors of the immune response that bind to cells expressing PD-1 to decrease their capacity for cytotoxic response. PD-L1 is normally expressed on dendritic cell (DC) wherein interactions between DC and T-cell via the PD1/ PD-L1 axis suppresses proliferation and potentiates apoptosis of T-cell (38, 39). Cancer cells can also express PD-L1 (40) to evade the cytotoxic effect of CD8+ T cell by suppressing cell proliferation through the PD-1/ PD-L1 axis (35, 41, 42). Our study employs anti-PD-1 antibodies to inhibit the suppressor effect of PD-1 expression by cancer cells. Consistent with this reasoning, there are reportes that blocking the PD-1/PD-L1 axis lead to increased expression of granzyme B in CD8+ T-cells in the presence of antigen along with increased proliferation and anti-tumor function (27,39). In our previous study, we reported the synergistic effect of aFP and anti-PD-1 inhibitors leading to the increase of TAA-specific CD8+ T-cells systemically (11). In our present study, we confirmed that aFP + anti-PD-1 therapy led to an increase in XCR1+ DC in drainage lymph nodes, further validating the mechanism by which the two interventions augment anti-cancer immunity. However, despite our observation of TAA-specific CD8+ T-cells in contralateral, untreated tumors with aFP + anti-PD-1 therapy, only 33% of untreated contralateral tumors achieved complete remission (11). We posited that the greater effect on local tumors over the abscopal effect on remote cancer cells was due to the higher ratio of CD8+ T-cells compared to the baseline T_reg_ cell population in the local tumor since the ratio in the untreated contralateral tumors was low and there was no significant difference compared to the no treatment control group.

To address this, we added a booster of the immune checkpoint agonist OX40 to increase the concentration of CD8+ T-cell systemically. OX40 is a secondary co-stimulatory immune checkpoint molecule, predominantly expressed by T-cells during antigen-specific priming in the presence of inflammatory cytokines such as Interleukin-1 (Il-1) (28). OX40’s ligand, known as OX40L or CD252, is expressed on activated professional antigen-presenting cells like dendritic cells (DCs), macrophages, and B cells (43, 44). Costimulatory signals via the OX40/OX40L axis promotes T-cell expansion, proliferation (12–14), and enhances the activation and effector function of killer T- and helper T-cells (15–17). Furthermore, there is some evidence that OX40 agonists may also be implicated in depleting the number of Foxp3+ T_reg_ cells, further contributing to the increased CD8+ T-cell to T_reg_ ratio (18–21). In this study, although T_reg_ depletion was not observed, triple therapy with aFP + anti-PD-1 + OX40 agonist led to a significant increase in CD8+ T-cells and complete remission was observed in both local and untreated contralateral tumors when treatment was started 6 days following tumor inoculation.

We propose the following mechanism for the increased efficacy of triple. 1) aFP-induced thermal denaturation of tissue releases TAA (11) and DAMPs (34), along with chemokines and cytokines, upregulating the expression of HSP70, HSP90, CALR and increasing expression of antigen on cell surfaces. 2) The chemokines, cytokines, and DAMPs recruit innate immune cells such as DCs (45, 46) which present TAA to T-cells via the interaction of MHC and TCR. In the presence of Il-1, TCR signaling stimulates the expression of OX40 on T-cells (Supplemental table 1)(28). 3) The presence of OX40 agonist induces T-cell proliferation, activation, and effector function leading to an increased number of granzyme B+ CD8+ T-cells. 4) The addition of PD-1 inhibitors blocks the PD-1/PD-L1 axis that would otherwise inactivate CD8+ T-cells and inhibit the immune response. Taken together, triple therapy works to enhance the immune response to antigen-bearing tumor cells, increasing the effector CD8+ T-cells systemically, leading to the effective eradication of cancer. In support of this mechanism, neither aFP + OX40 agonist nor aFP + anti-PD-1 were sufficient relatively to create a systemic immune response leading to effective tumor regression. In the aFP + OX40 agonist group, tumors could not be cured effectively and CD8+ T-cells did not infiltrate by 5 days after treatment started. There are reports that PD-1 is expressed on innate cells such as DCs and macrophage (38), and expression of PD-L1 by tumor cells is induced by exposure to Type I and II interferons (35, 47–49), and PD-L1 can even be secreted by tumor cells (50–52). Moreover, activating CD8+ T cells in tumors is reduced in the absence of blocking the PD-1/ PD-L1 axis (39). In our study, expression of PD-L1 on tumor cells and induction of interferon beta one at the RNA level in peripheral tumor microenvironment were observed after aFP treatment (Supplement figure 2 and table 1).

Therefore, we expect high PD-L1 expression on tumor cells in the presence of interferon beta induced by aFP, secretion of PD-L1 from tumor cells and the suppressive state of innate immunity at peripheral tumor microenvironment in the absence of PD-1 inhibitors leading to fewer DC, the interaction of MHC/ TCR between the innate and adaptive immune cells, and the infiltration of CD8+ T cells into tumors, leading to fewer cured mice than in triple therapy.

Similarly, in the aFP + PD-1 group, while we did see an increase in DCs over control, the population of CD8+ T-cells was unable to expand sufficiently in the absence of OX40 agonist to achieve the same robust increase in survival as triple therapy. Interestingly, therapy with anti-PD-1+ OX40 agonist alone led to complete tumor remission in 83% of mice compared to 100% in the triple therapy group when treatment was initiated 6 days after tumor inoculation. However, tumor volume curves show that the tumor volume peak in cured mice occurs two days later in the anti-PD-1+ OX40 agonist group when compared to the triple therapy group, indicating that the tumor shrank at a slower rate. We posit that the baseline excretion of TAA and expression of HSP70, HSP90 and CALR on tumor cells, plus the low expression of OX40 on T-cells in the absence of inflammatory cytokines, were insufficient to induce anti-tumor immunity. This is consistent with the flow cytometric data showing that total CD8+ T-cell populations in lymph nodes and tumors in the anti-PD-1+ OX40 agonist alone group were not increased compared to control 5 days after treatment. When treatment was started 12 days after tumor inoculation, only 30% of mice receiving anti-PD-1+ OX40 agonist alone were cured compared to 80% in the triple therapy group. We posit that the significant difference between experimental groups relates to the rate at which CD8+ T-cells proliferated. It is generally thought that initiating an immune response against larger tumors is difficult because they grow quickly and implement immune invasion techniques like recruiting T_reg_ and myeloid derived suppressor cells (MDSC), secreting immune suppressor cytokines, and down-regulating MHC class I molecules to reduce the expression of immunogenic antigens (53, 54). By delaying the onset of treatment, the proliferation of CD8+ T-cells in the anti-PD-1+ OX40 agonist alone group was not sufficient to produce a high enough concentration of T-cells to cure a rapid growing tumor. In terms of monotherapy with anti-PD-1 or OX40 agonist, neither increased the CD8+ or granzyme B+ CD8+ T-cell populations due to the absence of anti-PD-1 inhibitor or OX40 agonist, whose mechanism is mentioned above, resulting in poor survival outcomes.

In conclusion, the addition of an OX40 agonist to aFP and an anti-PD-1inhibitor effectively boosted systemic immunity. These effects are likely mediated by an immune cascade including the release of TAA by aFP treatment, increased proliferation of TAA specific CD8+ T cells induced by OX40 agonist, and inhibition of the immune suppressing PD-1/PD-L1 axis by anti-PD-1 inhibitor.

## Materials and Methods

### Cell lines

CT26WT murine colon and 4T1 murine mammary carcinoma cell lines (ATCC, Mannassas, VA) were cultured in RPMI Medium supplemented with 10% heat-inactivated fetal bovine serum, penicillin (100 U/mL), and streptomycin (100 mg/mL) (Sigma-Aldrich, Natick, MA) at 37° C in 5% CO_2_. Cultures were performed in 75 cm^2^ flasks (Falcon, Invitrogen, Carlsbad, CA).

### Animals

Six-week-old female BALB/c mice (Charles River Laboratories, Boston, MA) were used for the study. The care and handling of the animals were done in accordance with a protocol approved by the Subcommittee on Research Animal Care (IACUC) at Massachusetts General Hospital (MGH).

### Animal tumor model

In the primary experiment, mice were anesthetized by intraperitoneal injection of a ketamine (90 mg/kg) and xylazine (10mg/kg) cocktail before depilation of bilateral external thighs and subcutaneous inoculation with 3.5 × 10^5^ CT26WT cells. Anti-PD-1 blocking and/or anti-OX40 agonist antibodies (29F.1A12 and OX-86 respectively; BioXCell,West Lebanon, NH) were administered intraperitoneally at doses of 200 μg per mouse on days 6, 8, 10, 12, and 14 in the 6 day inoculation protocol and on days 12, 14, 16, 18 and 20 after tumor cell inoculation in the 12 day protocol (Figure 10). Tumor volume was determined at least 2 times per week by measuring the longest dimension and orthogonal dimension of the tumor with vernier calipers. Tumor volumes were calculated according to the formula volume = 4π/3×[(a+b)/4]^3^, where a and b represent the long and short axis lengths, respectively. If tumor volume exceeded 500 mm^3^ or showed severe ulceration, mice were removed from the study and defined as having reached their endpoint. In the T-cell depletion experiment, anti-CD8+ depletion antibodies (2.43; BioXCell,West Lebanon, NH) were administered intraperitoneally at a dose of 200 μg per mouse every 3 days from one day before tumor inoculation to removal of mice as an endpoint.

**Figure 10.**
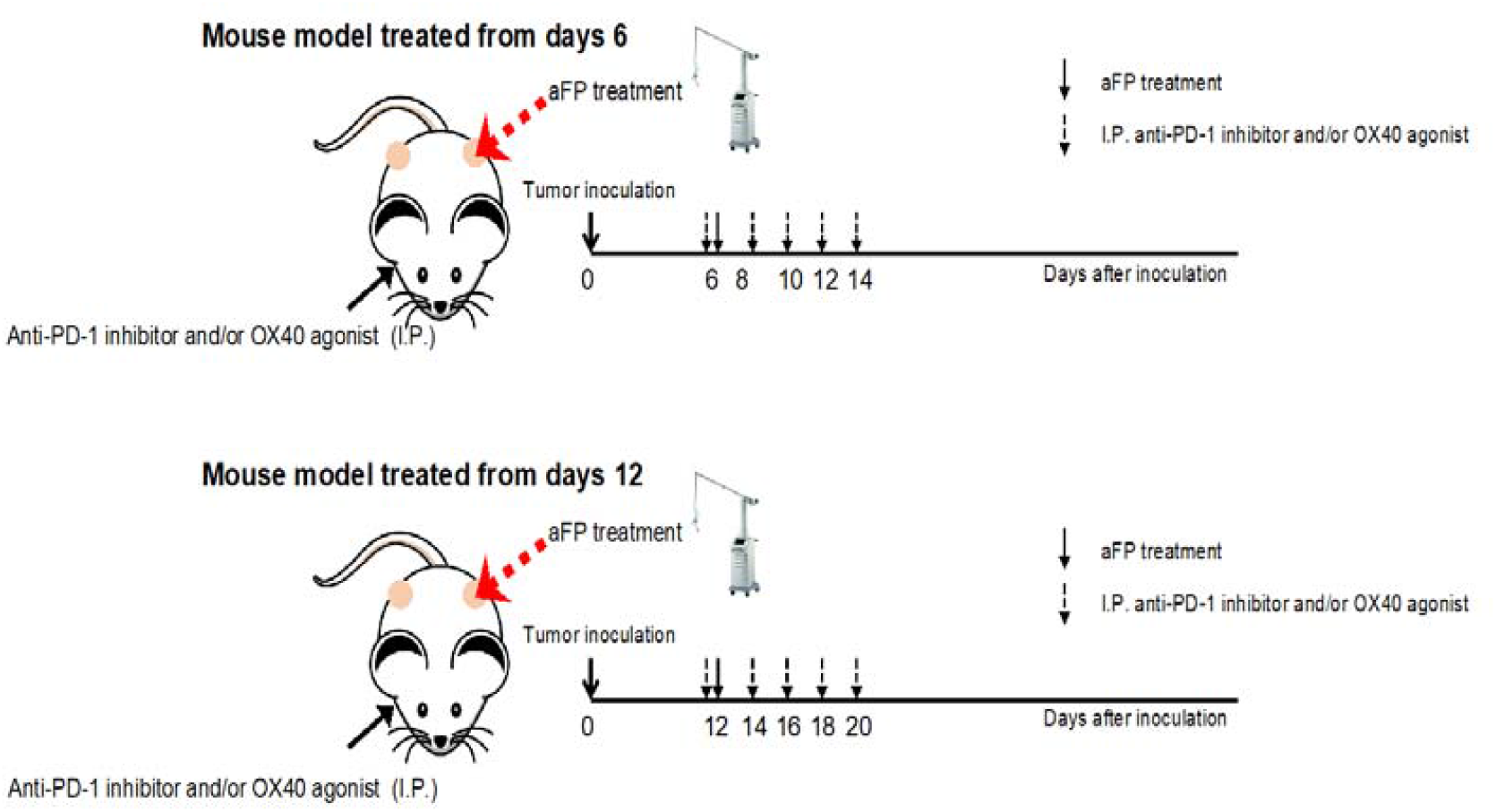
Experimental design scheme in the two-tumor model.

### Fractional CO_2_ laser irradiation

Ablative fractional laser treatment was performed on the tumor site on days 6 or 12 after tumor cell inoculation (Figure 10). 6 days post inoculation tumor size averaged 4 mm and measured an average of 7 mm at day 12. Exposures were performed on day 6 or day 12 with an Ultrapulse Encore CO_2_ laser (Lumenis Inc, Yokneam, Israel). A single pass was performed within a treatment area of 5×5 mm, with a pulse energy of 100 mJ, 5 % density, and 300 Hz frequency. No skin cooling was applied and the anesthesia was performed by intraperitoneal injection of a ketamine (90 mg/kg) and xylazine (10mg/kg) cocktail.

### Rechallenge

Mice surviving 90 days after tumor inoculation were rechallenged with matched tumor cells (3.5 × 10^5^ CT26WT) on the contralateral (right) leg from the previously aFP-treated side. Age-matched naive mice were inoculated with the same number of the cells in the right leg as the controls. Inoculated mice were monitored for another 60 days to confirm tumorigenesis.

### Flow cytometry analysis

Flow cytometry was used to analyze changes in CD3+ and CD8+ lymphocytes, regulatory T cell (T_reg_), and dendritic cell (DCs) populations in treatment and control groups. Freshly excised CT26WT tumors were mechanically dissociated through a 70 μm strainer into 6-well culture plates containing DNaseI (10 μg/ml, Roche; Nutley, NJ) and collagenase (10 mg/ml, Life technologies) and incubated for 60 minutes at 37°C. Dissociated cells were blocked using anti-CD16/CD32 antibody (93) for 15 minutes at 4°C before being stained with anti-CD45 (30-F11), -CD3 (145-2C11), -CD4 (RM4-5), -CD8a (53-6.7), -CD103 (2E7), -OX40 (OX-86; all from eBioscience; Santa Clare, CA), -CD11c (HL3; BD Biosciences; San Jose, CA), -IA/IE (M5/11415.2), -CCR7 (4B12), or -XCR1 (ZET; all from BioLegend; San Diego; CA) antibodies for 30 minutes at 4°C. Following staining for surface markers, cells were fixed and permeabilized for 30 minutes at room temperature using the Foxp3/Transcription Factor Staining Buffer Set (eBioscience) according to the manufacturer’s instructions. Permeabilized cells were incubated overnight at 4°C with anti-Foxp3 (FJK-16s), -Ki67(SolA15), or -granzyme-B (NGZB; all from eBioscience) antibodies. Flow cytometry was performed the day after staining with a Fortessa X-20 (BD Biosciences).

### Statistics

All experiments were repeated at least once. All statics analyses were performed with GraphPad Prism 9.0 (GraphPad Software). All values are expressed as the mean ± SD. Flow cytometric results were compared with one-way ANOVA. Survival analysis was performed using the Kaplan-Meier method and a log-rank test. Values of P < 0.05 were considered statistically significant.

## Acknowledgments

The authors wish to thank Lumenis for providing the CO2 laser as a gift, and Tuanlian Luo for preparing H & E stained slides.

## Author contribution

M.K. contributed to the conception, and design of the work and the acquisition, analysis, and interpretation of data; J.G. contributed to the acquisition of data and the proofreading of the manuscript; Z.T. contributed to the acquisition of data; S.D. contributed to the interpretation of data; D.M. contributed to the conception of the work.

## Additional Information

### Competing financial interests

Patent application number 63/088,281 has been filed (M.K. and D.M.) based on this work (“Intratumoral and Systemic Immunization Using Fractional Damage-Creating Device With Checkpoint Molecules for Cancer Therapy”)

## Figure and Tables

**Supplementary figure 1.**
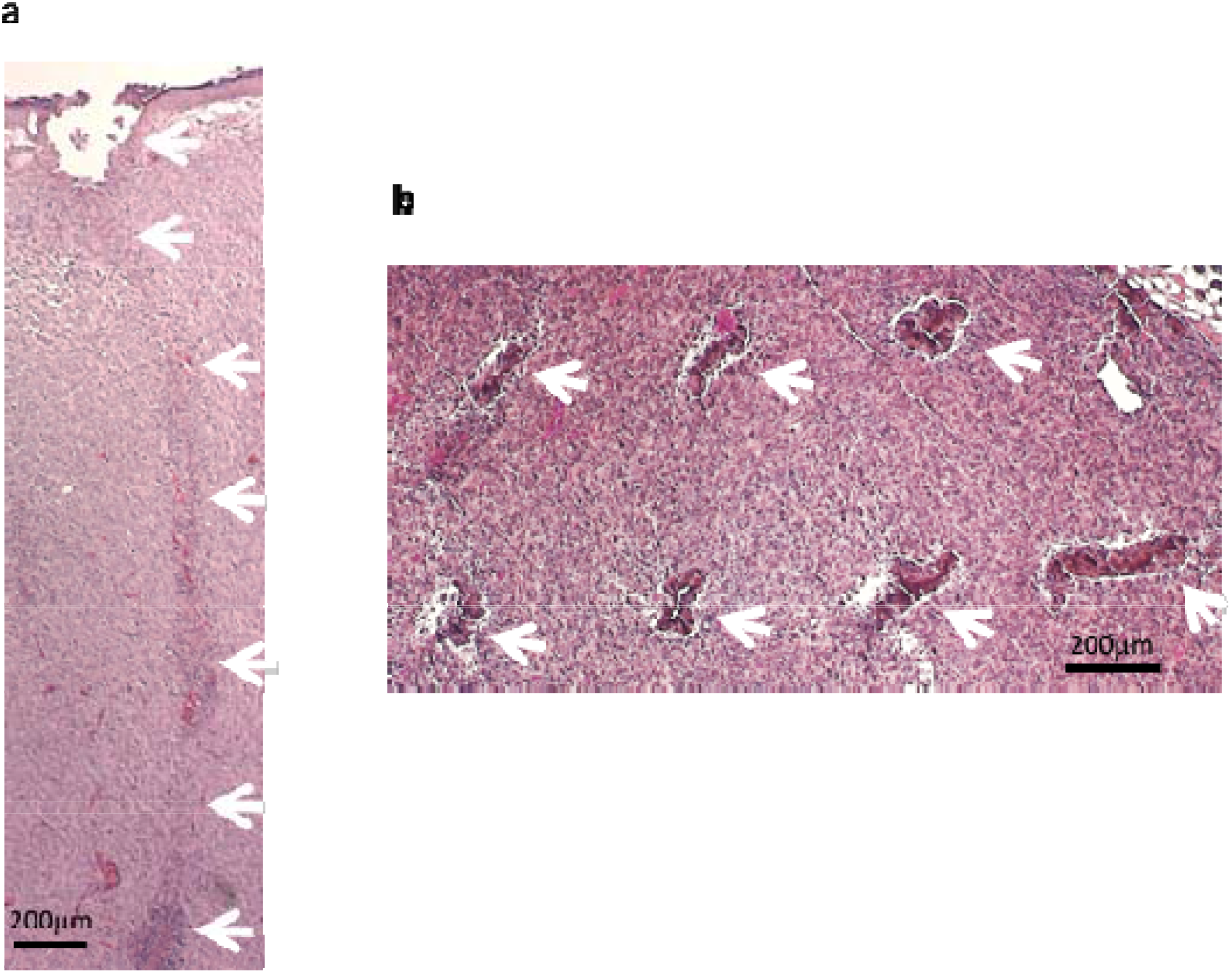
H & E stained histologic figures of tumor tissue harvested immediately after ablative fractional photothermolysis (aFP) laser treatment with a pulse energy of 100mJ at a nominal density of 5 %. (a) Parallel view to trajectory of laser beam (b) Orthogonal view to trajectory of laser beam. White arrows indicate an ablated hole characteristic of aFP procedures. The ablated hole appeared to be collapsed and distorted within the tumor tissue.

**Supplementary figure 2.**
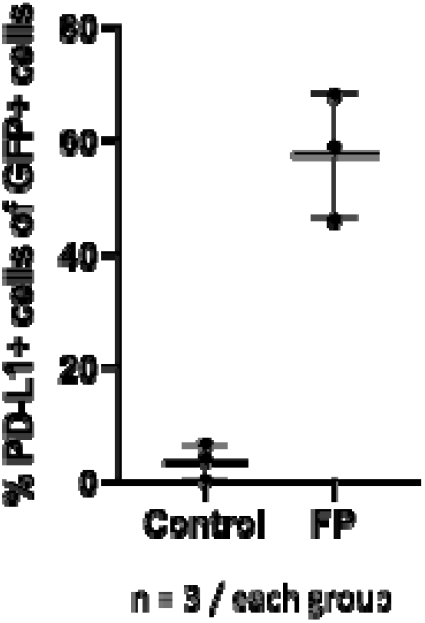
To investigate the up-regulation of PD-L1 receptors in tumors after aFP treatment, Green fluorescence Protein (GFP) gene was transfected to CT26WT cells using a lentivirus vector (GenTarget Inc). Both aFP-irradiated and non-irradiated tumors were harvested 3 days after irradiation. For isolation of GFP+ CT26WT cells, fresh tumors were dissociated mechanically filtering through 70 μm strainer on 6 well culture plate with DNaseI (10 μg/ml, Roche; Nutley, NJ) and collagenase (10 mg/ml, Life technologies) and incubated 60 minutes at 37°C. The dissociated cells were stained with anti-PD-L1(MIH7, BioLegend; San Diego; CA)) for 30 minutes at 4°C. Following staining for surface markers, cells were fixed by Foxp3/ Transcription Factor Staining Buffer Set (eBioscience; Santa Clare, CA) according to the manufacturer’s instructions at room temperature for 30 minutes. After fixation cells were suspended with washing buffer at 4°C over night. The next day the stained cells were analyzed on Fortessa X-20 (BD Biosciences; San Jose, CA).

**Supplementary table 1.** Data of RNA Sequence (aFP vs control group) To investigate the up-regulation of IL-1 and IFN-beta at the RNA level, we measured RNA expression of IL-1 and IFN-beta in tumors using RNA sequence analysis. aFP-irradiated tumor and non-irradiated tumor were harvested 24 hours after irradiation. RNA was extracted using RNeasy mini kit (Qiagen) and RNA sequencing was performed by a commercial company (Beijing Genomics Institute). Each group contain 3 tumors. The table shows aFP-treatment induces IL-1 and IFN-beta expression in tumor cells.

| Symbol | log2FoldChange(aFP/Control) | Probability | Up/Down-Regulation(aFP/Control) |
| --- | --- | --- | --- |
| Hspa1a | 7.725630695 | 1 | Up |
| Hspa1b | 8.309049618 | 0.99999999 | Up |
| Il1a | 3.746162754 | 0.99253465 | Up |
| Il1b | 3.446302695 | 0.999999807 | Up |
| Ifnb1 | 4.009460329 | 0.987913968 | Up |

## Notes

**Conflict of interest statement:** Patent application number 63/088,281 has been filed (M.K. and D.M.) based on this work (“Intratumoral and Systemic Immunization Using Fractional Damage-Creating Device With Checkpoint Molecules for Cancer Therapy”)

### Competing Interest Statement

Conflict of interest statement: Patent application number 63/088,281 has been filed (M.K. and D.M.) based on this work (Intratumoral and Systemic Immunization Using Fractional Damage-Creating Device With Checkpoint Molecules for Cancer Therapy)

### Summary of Updates

Added Zhipeng Tao's contribution in the section of author contribution

